# Mutational processes impact the evolution of anti-EGFR antibody resistance in colorectal cancer

**DOI:** 10.1101/2020.05.07.082339

**Authors:** Andrew Woolston, Louise J Barber, Beatrice Griffiths, Nik Matthews, Sheela Rao, David Watkins, Ian Chau, Naureen Starling, David Cunningham, Marco Gerlinger

**Affiliations:** Translational Oncogenomics Laboratory, The Institute of Cancer Research, London SW3 6JB, UK; Tumour Profiling Unit, The Institute of Cancer Research, London SW3 6JB, UK; Gastrointestinal Cancer Unit, The Royal Marsden Hospital, London SW3 6JJ, UK

**Keywords:** cancer evolution, drug resistance mechanisms, drug-induced mutagenesis, colorectal cancer, cetuximab, EGFR antibodies, mutational signatures, signature 17, predictive biomarkers

## Abstract

Anti-EGFR antibodies such as cetuximab are active against *KRAS/NRAS* wild-type colorectal cancers (CRC) but acquired resistance invariably evolves. Which mutational mechanisms enable resistance evolution and whether adaptive mutagenesis, a transient cetuximab-induced increase in mutagenesis, contributes in patients is unknown. We investigated this in exome sequencing data of 42 baseline and progression biopsies from cetuximab treated CRCs. Mutation loads did not increase from baseline to progression. Evidence for a contribution of cetuximab-induced mutagenesis was limited. However, the mutational Signature 17 was a key contributer of specific *KRAS/NRAS* and *EGFR* driver mutations that are common at acquired resistance. Signature 17 activity before treatment predicted shorter progression free survival. This demonstrates the utility of mutational signatures to predict cancer drug resistance evolution.

**SIGNIFICANCE:** Drug resistance evolution occurs ubiquitously in solid tumours during treatment with targeted drugs. Biomarkers that can be assessed prior to treatment to predict the time to resistance evolution and the genetic resistance mechanisms that will evolve have not been described. We identified the mutational Signature 17 as the first candidate biomarker that predicts shorter time to progression and several specific *KRAS/NRAS* and *EGFR* mutations that will likely evolve in CRCs during cetuximab treatment. Understanding the mutational mechanism underlying Signature 17 may open opportunities to delay resistance acquisition. The potential of mutational signatures to predict resistance to a broader range of drugs in other tumor types should be assessed.

## INTRODUCTION

The anti-EGFR antibody (EGFR-AB) cetuximab is active against many *KRAS/NRAS* wild-type metastatic colorectal cancers (CRCs) (Karapetis et al., 2008; Van Cutsem et al., 2015). However, resistance invariably evolves within several months. Darwinian selection of subclones that harbor mutations in *KRAS, NRAS* and *EGFR* is among the commonest mechanisms of acquired resistance (Bettegowda et al., 2014; Misale et al., 2012; Montagut et al., 2012; Woolston et al., 2019). Pre-treatment biomarkers that can predict the time to resistance evolution and the specific resistance mechanism that will evolve have not been identified (Lipinski et al., 2016; Maley et al., 2017).

Mutation generation is central to resistance evolution, and mutational signature analysis can be used to dissect cancer mutational processes (Alexandrov et al., 2013; Gerlinger and Swanton, 2010). Yet, how the activity of specific mutational signatures enables or constrains the evolution of cetuximab resistance in CRCs is unknown. Resistance evolution may furthermore be influenced by the timing of specific mutational processes. The pre-existing drug resistance model assumes that such mutations are already present in small subclones before EGFR-AB exposure, making the evolution of acquired resistance inevitable (Figure 1A) (Diaz et al., 2012). Recently, a model of ‘adaptive mutagenesis’ has been proposed in which cetuximab treatment triggers a transient down-regulation of mismatch repair (MMR) and homologous recombination (HR) DNA repair proteins and increased expression of low-fidelity DNA polymerases, which together promote mutation generation in CRC cells (Russo et al., 2019). Such drug-induced mutagenesis could increase the probability of resistance mutation acquisition *during* treatment (Figure 1A). Importantly, these are preclinical observations and it is unknown whether cetuximab-induced mutagenesis is relevant in patients (Gerlinger, 2019). More generally, it remains undetermined whether any specific mutational signatures change through cetuximab treatment and which signatures generate the majority of resistance mutations observed in the clinic.

**Figure 1:**
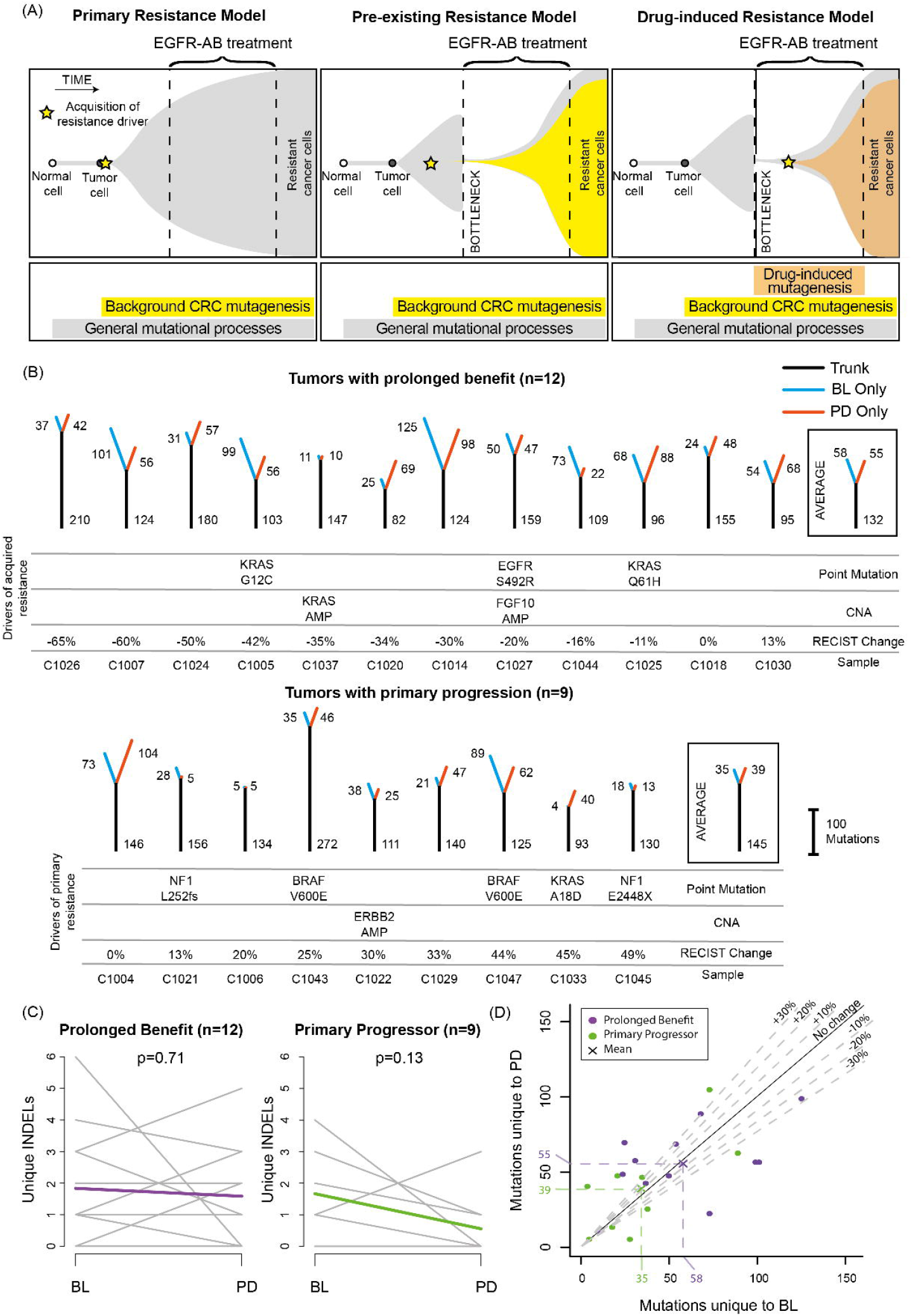
Cetuximab resistance models and analysis of mutation loads in 21 tumors treated with single-agent cetuximab. (A) Models of primary and acquired cetuximab resistance and their relationship to mutation signature activity. (B) Mutation trees for 21 tumors from the Prospect-C trial. Grouped into cases with prolonged benefit and primary progression. Numbers next to the trunk or the branches indicate the number of somatic mutations. Cetuximab resistance driver mutations and copy number aberrations (CNA) identified in (Woolston et al., 2019) are shown. The RECIST change indicates the change of the sum of radiological tumor measurements based on RECIST criteria from BL to the time of best response. (C) Change of the unique INDEL numbers from BL to PD. Colored lines show the mean. The p-values were calculated with a paired t-test. (D) Unique mutation loads for each tumor at BL versus PD. The dashed lines indicate a relative increase or decrease by 10%, 20% or 30%.

Our aim was to assess the activity of distinct mutational mechanisms in serial biopsies from initially *KRAS/NRAS* wild-type CRC patients who were treated with single-agent cetuximab in a prospective biomarker discovery trial. Drug treatment forces the cancer cell population through an evolutionary bottleneck. We reasoned that this should reveal the mutational signatures operating before or during drug treatment as these become increasingly clonal and hence detectable by exome sequencing. Cetuximab-induced mutagenesis should increase both, mutation loads and the specific mutational signatures that are characteristic of these mechanisms in patients who benefit (Figure 1A). In contrast, no changes in mutation loads and signatures would be expected in patients with primary progression where cetuximab lacks activity. We furthermore questioned which mutational mechanisms are most relevant for the generation of the hotspot driver mutations that most frequently evolve at acquired resistance.

## RESULTS

### Clinical trial samples

The patient characteristics and biopsy analysis of the Prospect-C phase II trial have been described previously (Woolston et al., 2019). In brief, biopsies were taken at baseline (BL) before cetuximab initiation and a second biopsy at progressive disease (PD) from *KRAS/NRAS* wild-type CRCs. 42 paired BL and PD biopsies from 21 patients were successfully analyzed by exome sequencing and had similar cancer cell contents. No tumor showed evidence for MMR deficiency at BL (Woolston et al., 2019). Progression at or before the first per-protocol CT scan (scheduled at 12 weeks) had been classified as ‘primary progression’ (n=9). The remaining tumors were considered to have obtained ‘prolonged benefit’ (n=12) from treatment.

### Temporal change of mutation loads

Mutation trees were generated to analyze the evolutionary relationship of cancer cells in BL and PD biopsies and changes in the mutation load (Figure 1B). The trunk represents mutations present in both samples whereas branches indicate mutations unique to BL or PD samples. Truncal mutation loads were similar between tumors with prolonged benefit and primary progression (p=0.53, t-test). Cancers with prolonged benefit had higher unique mutation numbers compared to primary progressors (mean sum of BL and PD: 113 and 73, respectively, p=0.11, t-test). Although this was not significant, it likely indicates a cetuximab-induced population bottleneck that diminishes treatment-sensitive subclones which are replaced by subclones with distinct mutations at acquired resistance, whereas subclones at BL and PD are more similar in primary progressors.

The number of unique mutations did not significantly change from BL to PD, neither in tumors with prolonged benefit (p=0.74) nor in those with primary progression (p=0.62, paired t-test). The number of unique small insertions and deletions (INDELs), which are a characteristic of MMR-deficiency (Kim et al., 2013), also showed no increase from BL to PD (prolonged benefit: p=0.71; primary progression: p=0.13, paired t-test) (Figure 1C).

The absence of a population bottleneck in primary progressors is a potential source of bias as these may harbor higher numbers of subclones at PD, leading to higher subclonal mutation loads than in tumors with prolonged benefit where subclones were pruned. We therefore repeated the analysis by only considering clonal mutations in each sample. This showed a small but non-significant increase of unique mutations in tumors with prolonged benefit (+11.0%, p=0.66) but an even larger increase in primary progressors (+48.8% p=0.20, paired t-test, Figure S1).

We furthermore considered that cetuximab-induced mutagenesis may only be active in a subgroup of tumors. 5/12 (41.7%) cases with prolonged benefit showed a ≥20% increase of the unique mutation load at PD but also 4/9 (44.4%) of tumors with primary progression (Figure 1D). Thus, although mutations can increase in individual tumors after treatment, this fraction did not differ between these groups.

Taken together, we found no evidence for a rise in the mutation load through cetuximab treatment. This mirrors results from Russo et al. (Russo et al., 2019) who described only a negligible change in mutation burden in cetuximab treated CRC cell lines analyzed by exome sequencing. Exome sequencing only analyzes ∼1% of the genome which may be insufficient to reliably detect an increase of mutations across the genome. However, these results show that if drug-induced mutagenesis is active, the impact on the mutation load in the protein-coding genome is small.

### Microsatellite tract length variability

Cetuximab-induced mutagenesis increased the accumulation of INDELs in microsatellite tracts in CRC cell lines (Russo et al., 2019). Applying the same approach to assess the length variability of microsatellite tracts to the Prospect-C trial cohort showed no increase from BL to PD in tumors with prolonged benefit or with primary progression (Figure 2). Restricting the analysis to those tumors with ≥20% increase in the unique mutation load at PD also showed no change. Thus, we found no evidence for a cetuximab-induced increase in microsatellite tract length variability.

**Figure 2:**
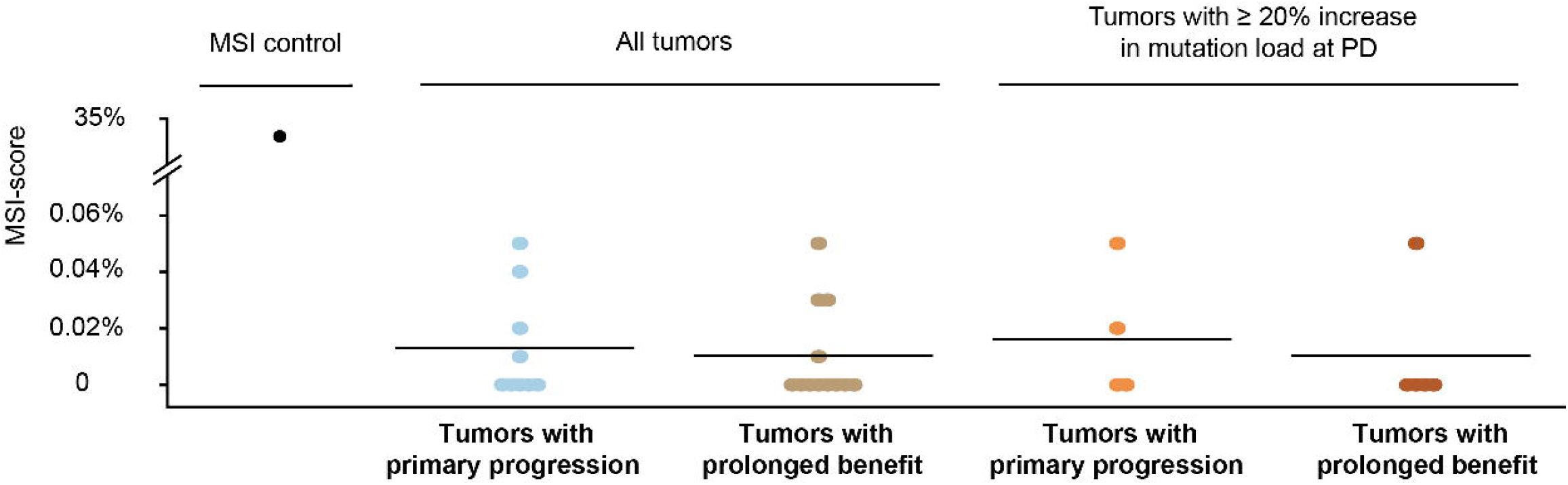
Microsatellite length variability analysis with the MSIsensor algorithm. MSI-scores indicate the percentage of microsatellite and homopolymer loci with an increased read length variability at PD compared to BL. Horizontal bars show the mean MSI-score for each group. The MSI-score of the only MMR-deficient tumor from the Prospect-C trial (which has not been included in any other analyses as no paired PD sample was available) in comparison to the matched blood sample is shown as a control for correct MSI detection.

### Temporal changes of mutational signatures

Mutational signature analysis (Alexandrov et al., 2013) should reveal changes in the activity of mutagenic processes independent of mutation loads. All single nucleotide substitutions and the two flanking bases were analyzed, corresponding to 96 tri-nucleotide sequence motifs. To limit the impact of signature bleeding, which can lead to the misassignment of mutations to signatures with high similarity (Maura et al., 2019), we only included signatures that have previously been described in CRC (Signatures 1 and 5 which are active in most cancer types (Alexandrov et al., 2015), Signature 6 which is typical for CRCs with MMR-deficiency (Sveen et al., 2017), Signature 17 which has been reported in chemotherapy treated CRCs (Christensen et al., 2019; Pich et al., 2019)) and signatures that may increase through cetuximab-induced mutagenesis (the HR-deficiency Signature 3 and additional MMR-deficiency Signatures 15, 20 and 26 (Alexandrov et al., 2013; Meier et al., 2018)).

Individual tri-nucleotide motifs only showed small changes from BL to PD (Figure 3A). After assigning these to mutational signatures, Signature 1 and signatures with a broad range of substitution motifs (Signature 3, Signature 5) were most abundant (Figure 3B and C). Signature 17, which is characterized by a strong predominance of T>G/T>C mutations in a CTT context, was assigned the next highest number of mutations. Compared to Signatures 1 and 3 which were active in most samples, Signatures 5 and 17 were only active in a subset (52% and 62%, respectively). Only Signature 3 showed an increase from BL to PD in the prolonged benefit group but this was small (+2%, mean: +2.5 mutations/tumor) and it also increased in primary progressors (Figure 3C).

**Figure 3:**
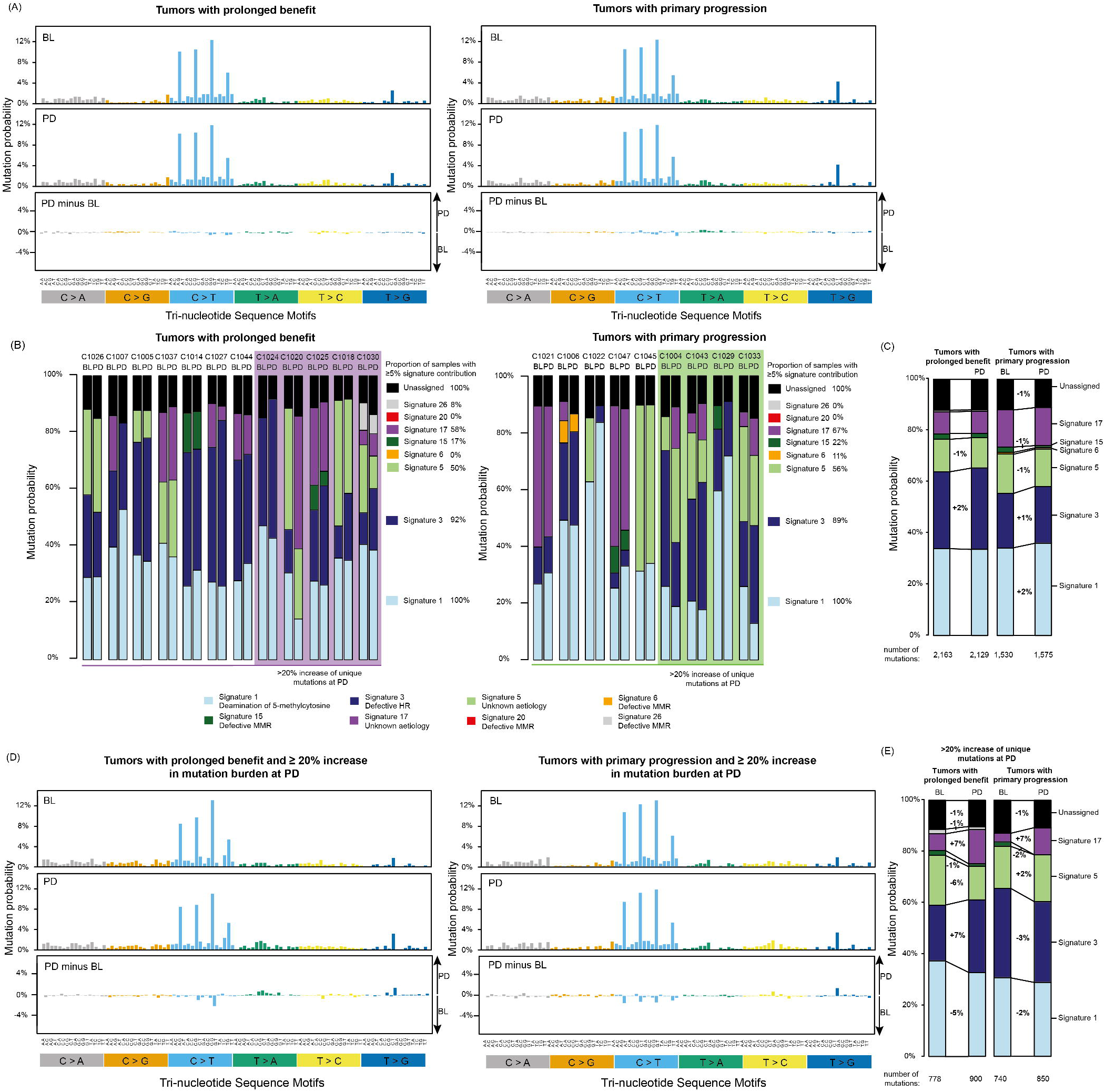
Mutational signatures in tumors treated with cetuximab. (A) 96 tri-nucleotide motif plot of all single base substitutions at prior to cetuximab treatment (BL) and at progression (PD). The bottom panel shows the difference between BL and PD. (B) Attribution of single base substitutions to COSMIC mutational signatures shows the contribution of each signature to individual samples at BL and PD. (C) COSMIC signature contribution for the combined group of cases with prolonged benefit or primary progression. (D) Mutational signatures in tumors where a ≥20% increase in unique mutation burden was found at PD. (E) COSMIC mutational signature contribution for the combined group of cases with prolonged benefit or primary progression which also showed a ≥20% increase in unique mutation burden.

Focusing only on the 5 tumors with prolonged benefit and a ≥20% increment in unique mutations also revealed the largest increase for Signature 3 (+7%, mean: +17.2 mutations/tumor, Figure 3D-E). Signature 17 also appeared to increase (+7%, mean: +13.6 mutations/case) but this was driven by a single tumor with a large increment (Figure 3B). Furthermore, signature 17 also rose by 7% in tumors with primary progression and a ≥20% mutation increase at PD. Thus, the mutational process that generates Signature 17 is unlikely to be promoted by cetuximab.

Taken together, the HR-deficiency Signature 3 was the only signature that noticeably increased at PD in the prolonged benefit group. This would be consistent with a cetuximab-induced loss of HR-fidelity (Russo et al., 2019). Yet, the mutational increment was small and an increase was only detected in some cases despite a median cetuximab treatment duration of 26 weeks (range: 18-96 weeks) in the prolonged benefit group.

### Signature 17 disproportionally contributes to driver mutations enriched at acquired resistance

*KRAS, NRAS* and *EGFR* mutations are the commonest mutational mechanisms of acquired cetuximab resistance in CRC (Bettegowda et al., 2014; Misale et al., 2012; Montagut et al., 2012; Woolston et al., 2019). Mutations in these genes at acquired resistance differ from those found in treatment-naïve CRCs: *EGFR* mutations at acquired resistance disrupt cetuximab binding epitopes and do not occur in untreated CRCs as they provide no fitness advantage in the absence of treatment (Arena et al., 2015). Furthermore, comparison of biopsy- and ctDNA-sequencing results of CRCs with acquired cetuximab resistance (Bettegowda et al., 2014; Khan et al., 2018; Woolston et al., 2019) to biopsy sequencing data of *KRAS/NRAS* mutant treatment-naïve CRCs (Cancer Genome Atlas Research et al., 2013) showed that *KRAS/NRAS* codon 12/13 mutations were 1.7-fold lower and codon 61 mutations 4.2-fold higher in tumors with acquired resistance compared to tumors with expected primary resistance. Q61H mutations showed the largest increase, by 11.8-fold (Figure 4A). Results were similar for the CORRECT trial that assessed *KRAS* mutations in ctDNA at acquired EGFR-AB resistance (Figure S2) (Tabernero et al., 2015). *KRAS* Q61H mutations were 21.1-fold higher than in treatment-naïve *KRAS* mutant CRCs. Based on our observation that signature contributions varied between tumors in the Prospect-C trial, we questioned whether signature activity before cetuximab initiation influences which resistance driver mutations evolve at acquired resistance.

**Figure 4:**
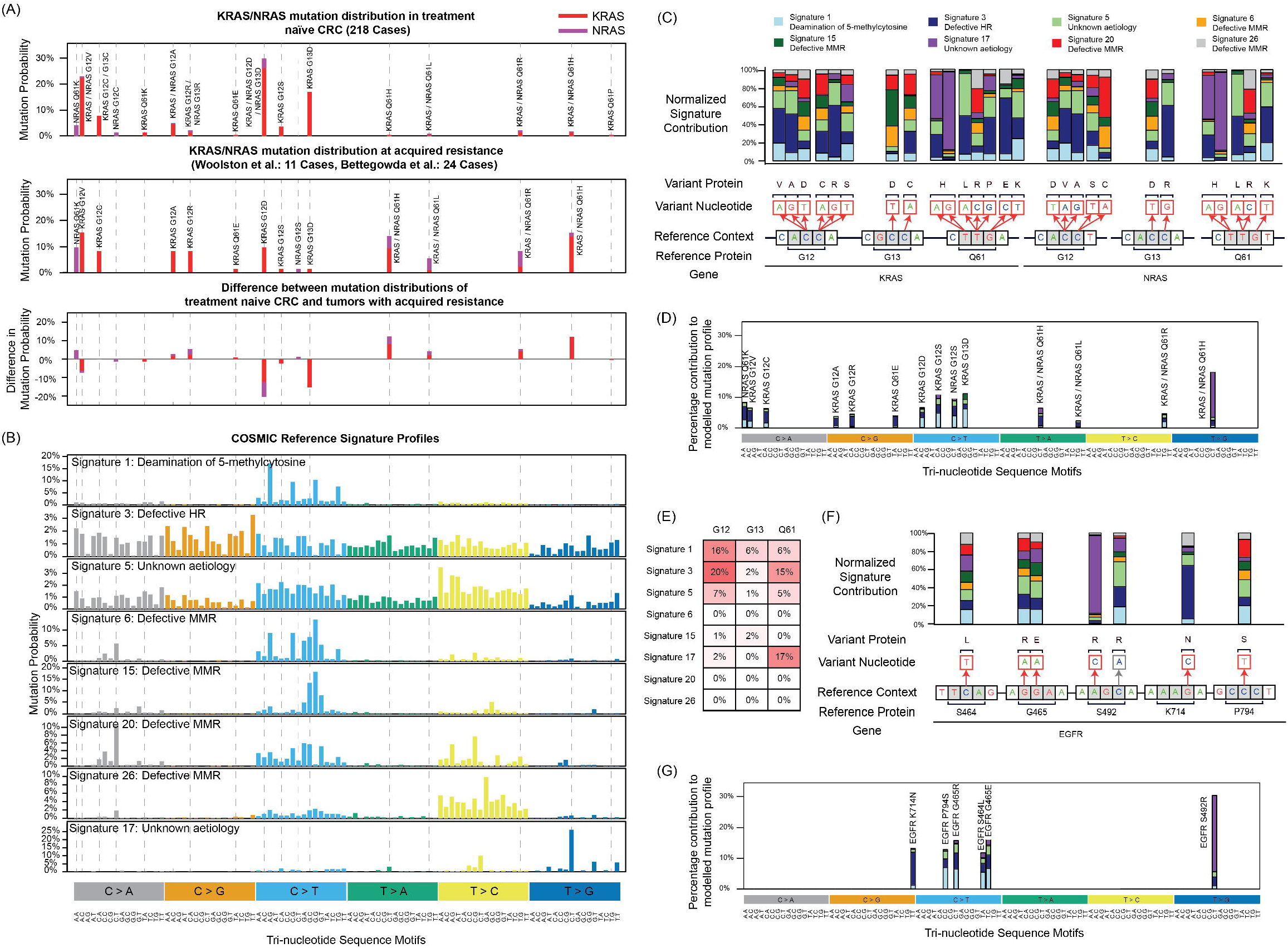
Relationship of mutational signatures to specific KRAS/NRAS and EGFR mutations. **(A)** KRAS/NRAS codon 12/13/61 mutation frequency in treatment naïve CRCs from the TCGA Pan-Cancer study versus those identified in CRCs with acquired EGFR-AB resistance(Bettegowda et al., 2014; Khan et al., 2018; Woolston et al., 2019). (B) Signature profiles of all COSMIC v2 mutational signatures included in the analyses of the Prospect-C cohort. (C) Relative contribution of each of the signatures in B to the tri-nucleotide sequence motifs corresponding to the indicated KRAS/NRAS mutations when an equal number of mutations is generated with each signature. All reference contexts in the figure show the main genomic strand. (D) Modelling of the relative contribution of each of the signatures in B across all indicated KRAS/NRAS mutations when the observed mutational signature distribution at BL in cases with prolonged benefit is taken into account. (E) Heatmap of the contribution of each signature in D to the mutations in KRAS/NRAS codons 12, 13 and 61. (F) Repeat of the analysis in panel C for EGFR mutations. (G) Repeat of the analysis in panel D for EGFR mutations.

We first compared *KRAS/NRAS* mutation profiles in CRC (Figure 4A) to the mutational signatures profiles (Figure 4B). The broad profiles of Signatures 3 and 5 overlapped with most hot-spot mutations. However, the remaining signatures only overlapped a few, indicating that the activity of these signatures could influence distinct evolutionary outcomes. We hence calculated the probability that each of the mutational signatures generates specific *KRAS/NRAS* mutations (Figure 4C). Intriguingly, Signature 17 showed a strong preference for generating *KRAS/NRAS* Q61H mutations and almost exclusively generated the T>G mutation that was most enriched at acquired cetuximab resistance. Thus, Signature 17 activity could critically influence the probability that these mutations evolve.

We therefore modelled the *KRAS/NRAS* mutation distribution that would be generated in prolonged benefit cases based on the observed signature contribution at BL (Figure S3, Figure 4D). Despite the higher activity of Signatures 1, 3 and 5, Signature 17 was the largest contributor to the generation of Q61H mutations (Figure 4E). In contrast, codon 12/13 mutations were predominantly generated by Signatures 1 and 3.

We similarly assessed how mutational signatures relate to *EGFR* mutations at acquired resistance (Figure 4F). The *EGFR* S492R A>C mutation is also almost exclusively generated by Signature 17. K714N shows a strong preference for generation by Signature 3. When the signature contributions at BL in tumors with prolonged benefit was taken into account, these two signatures are the main drivers generating these specific mutations (Figure 4G).

Together this indicates that Signature 17 activity is critical for the evolution of *KRAS/NRAS* Q61H and EGFR S492R A>C mutations at acquired resistance.

### Signature 17 activity as a predictor of mutation evolution and progression free survival

To substantiate the relevance of Signature 17 in patients, we investigated whether Signature 17 activity can predict the evolution of specific drivers at acquired resistance and of progression free survival (PFS) in the Prospect-C trial. Signature 17 was active at BL in five cases (Figure 5A). *KRAS/NRAS* Q61H mutations evolved in four of these and an *EGFR* S492R A>C mutation in one. No *KRAS/NRAS* Q61H or *EGFR* S492R mutations were identified in tumors without Signature 17 activity. This statistically significant enrichment suggests that Signature 17 activity canalizes the evolution of these resistance driver mutations. Furthermore, Signature 17 predicted for a shorter PFS in the prolonged benefit group but not in primary progressors (Figure 5B).

**Figure 5:**
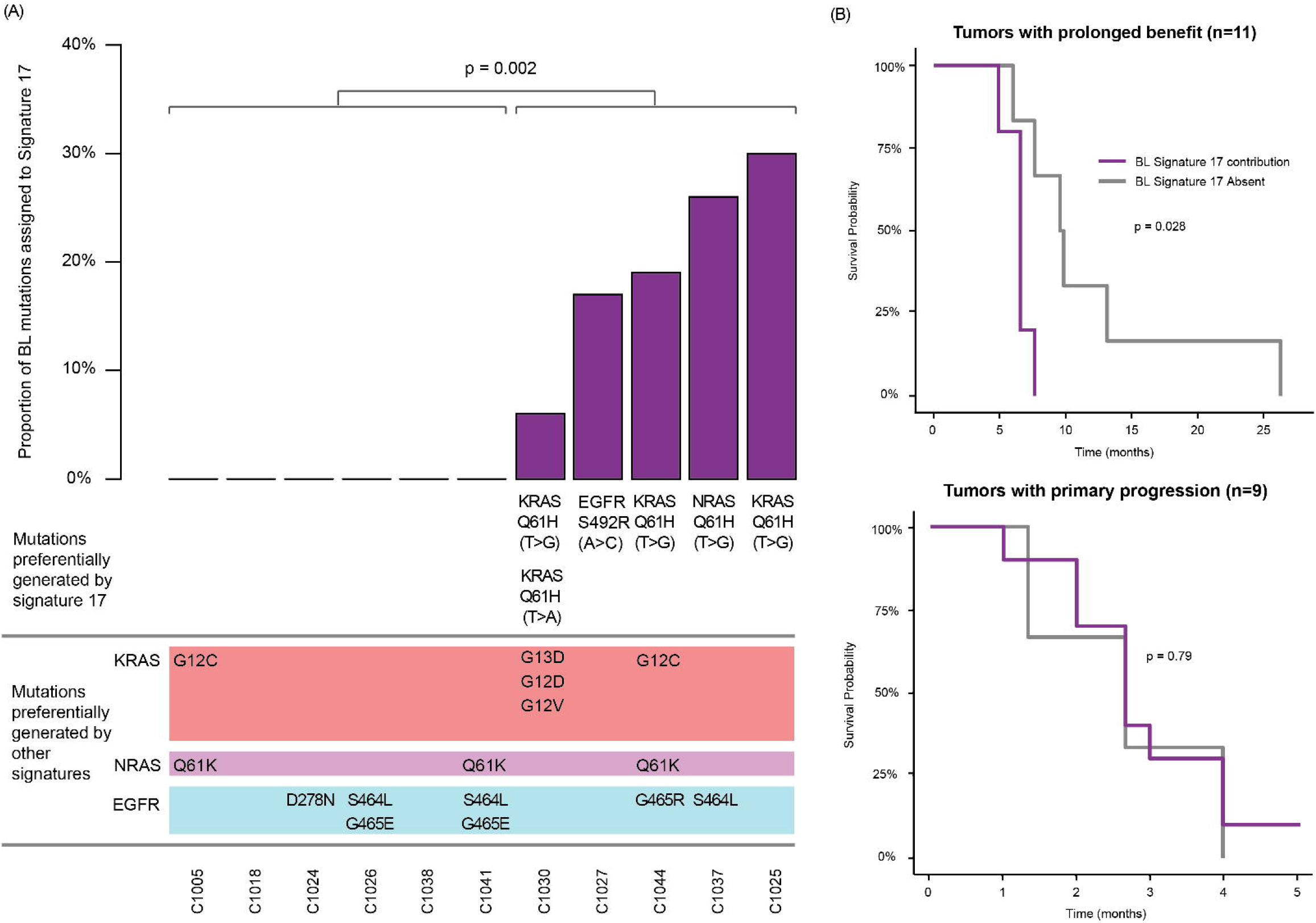
Association of Signature 17 at BL with specific KRAS/NRAS and EGFR mutation evolution at the time of acquired resistance and with progression free survival. (A) Signature 17 contribution calculated from whole exome mutation analysis of BL biopsies for all prolonged benefit cases with available ctDNA sequencing versus resistance driver mutations in KRAS/NRAS and EGFR that were detected at PD. Statistical significance was assessed with the Fisher’s exact test. (B) Kaplan-Meier analysis of progression free survival for tumors with and without Signature 17 contribution at BL. Statistical significance was assessed with the Log-rank test.

## DISCUSSION

This analysis of CRCs that were prospectively treated with cetuximab revealed how mutational processes shape resistance evolution. We showed that *KRAS/NRAS* codon 61 mutations are 11.8-fold to 21.1-fold more common at acquired resistance to EGFR-AB than in treatment-naïve *KRAS/NRAS* mutant CRCs. Prior work suggested that Q61 mutations have a higher oncogenic potential than codon 12/13 mutations when KRAS expression is low and that this explains the codon bias at acquired resistance (Ali et al., 2017) but there is little evidence to support lower KRAS/NRAS protein expression at acquired resistance. We have now shown that Q61H is predominantly generated by Signature 17 which is undetectable in most treatment-naïve CRCs but present in 62% of advanced CRCs treated with chemotherapy (Bettegowda et al., 2014; Khan et al., 2018; Woolston et al., 2019). This provides a compellingly simple explanation for the codon bias of *KRAS/NRAS* mutations between primary and acquired resistance. Moreover, Signature 17 activity may also explain the high prevalence of the S492R mutation amongst *EGFR* mutations (Montagut et al., 2018). Datasets for independent validation of these novel findings are not available in the public domain but our results are strengthened by the use of data from a prospective clinical trial which limits selection biases and by four independent lines of evidence: We showed that Signature 17 disproportionally contributes to *KRAS/NRAS* Q61H and *EGFR* S492R mutation generation. Secondly, the observed signature contribution in BL biopsies leads to *KRAS/NRAS* Q61H mutation frequencies similar to those observed at acquired resistance in published studies. Thirdly, we showed that the presence of Signature 17 at BL correlated with the evolution of *KRAS/NRAS* Q61H and *EGFR* S492R mutations in individual patients. Finally, PFS was shorter in patients where Signature 17 was detectable at BL, suggesting that this signature increases cancer evolvability during cetuximab treatment.

Signature 17 activity may therefore not only allow to predict that *KRAS/NRAS* Q61H and *EGFR* S492R will likely evolve in a patient but may also be the first evolutionary biomarker that can predict shorter PFS with cetuximab treatment. These results furthermore raise the question whether inhibiting the process generating Signature 17 mutations can delay cetuximab resistance. Whether this is feasible remains to be determined as these mechanisms remain poorly characterized. Recent studies made some progress by showing similarities with the mutation profile of samples treated with 5-fluorouracil (Christensen et al., 2019; Pich et al., 2019).

We furthermore assessed whether there was evidence for cetuximab-induced mutagenesis (Russo et al., 2019). We found no increase of mutation loads at acquired resistance, nor evidence for cetuximab-mediated MMR-deficiency as relevant mutational signatures, INDELs, and microsatellite lengths appeared unaffected by cetuximab. However, we detected a 7% increase in Signature 3 mutations in some tumors with prolonged benefit. This may be the consequence of reduced HR-fidelity through cetuximab-induced mutagenesis in a subgroup of cases. Overall, Signature 3 substantially contributed to *KRAS/NRAS* and *EGFR* mutations but as this signature was already active in most tumors at BL and only a small proportion of signature 3 mutations at PD can potentially be attributed to cetuximab treatment. Thus, despite the strong functional evidence for cetuximab-induced mutagenesis in CRC cell lines (Russo et al., 2019), our analysis in patients shows that its contribution to cetuximab resistance evolution and the impact on mutation loads in the protein coding genome are likely small.

There are limitations of our analysis. Although it is the largest series of biopsies from cetuximab treated CRCs that has been interrogated by exome sequencing, the analysis of further cohorts, ideally by whole-genome sequencing, may strengthen the evidence for drug-induced mutagenesis. Moreover, Signature 3 is a ‘broad’ signature with mutation motifs overlapping those of Signature 5 (Maura et al., 2019), which may lead to signature bleeding. Signature 3 was not observed in CRCs series that were analyzed by whole-genome sequencing (Alexandrov et al., 2013). Prior exposure of our CRC cases to chemotherapy and analysis of signatures from exome as opposed to whole-genome data may explain this.

Taken together, we provided proof of principle that mutation signatures can influence and predict the evolution of cetuximab resistance in CRC patients. This defines a novel strategy for the development of evolutionary biomarkers in precision cancer medicine.

## Acknowledgements

MG received funding from Cancer Research UK, the NIHR Biomedical Research Centre for Cancer at the Institute of Cancer Research and the Royal Marsden Hospital, Cancer Genetics UK, the Constance Travis Trust, the European Research Council (ERC) under the European Union’s Horizon 2020 research and innovation programme (grant agreement No. 820137). The paper is dedicated to the memory of Tim Morgan who supported this work with a generous donation.

## Author Contributions

AW and MG conceived and designed the study. MG funded and supervised the analysis. IC is the chief investigator of the Prospect C trial and obtained funding for the trial. SR, DW, NS, IC recruited treated patients in the Prospect C trial. LJB, BG, NM processed and performed genetic analysis of tumor samples. AW performed bioinformatics and statistical analysis. MG, AW and LJB wrote the manuscript and all authors approved of the final manuscript:

## METHODS

### Trial Design And Samples

Prospect-C is a single-arm phase II trial that investigated biomarkers of response or resistance to single-agent cetuximab in *KRAS/NRAS* wild-type metastatic CRCs. The trial has previously been described in detail (Woolston et al., 2019). The study was carried out in accordance with the Declaration of Helsinki and approved by the national UK ethics committee (approval number: 12/LO/0914). Written informed consent for trial participation and the molecular analysis of tumor biopsies was obtained from all patients.

### Somatic Mutation And Clonality Assessment

Published mutation calls were re-analyzed (Khan et al., 2018; Woolston et al., 2019). A mutation call with variant allele frequency (VAF) less than 5% was considered absent in either paired biopsy. The clonality of somatic variants was assessed as previously described (Woolston et al., 2019).

### Mutational Signature Analysis

The single base substitution mutation profile for each patient biopsy were fitted to the COSMIC (Tate et al., 2019) v2 signatures using *whichSignatures* in the deconstructSigs (Rosenthal et al., 2016) (v1.8.0) R library. The method was run with normalization option ‘exome2genome’ as recommended for whole exome data to scale the tri-nucleotide frequencies to the likely proportions observed across the genome. Signature contributions with weight <5% were discarded by specifying ‘signature.cutoff=0.05’. The error between the observed (tumor) and reconstructed (product) matrices was also included in the ‘unassigned’ signature contribution.

The mutation load assigned to individual signatures was calculated by multiplying the adjusted signature weights by the total single base substitutions for each biopsy. The mutation distribution that would be expected in *KRAS/NRAS* or *EGFR* genes based on the mutational signatures observed in the baseline biopsies from prolonged benefit cases was modelled by scaling the COSMIC signature profiles by the corresponding signature contribution.

### Microsatellite Tract Length Analysis

MSIsensor (Niu et al., 2014) (v0.6) *scan* was run on the complete hg19 reference sequence to identify homopolymer and microsatellite regions with a minimum of five consecutive repeats. This identified a total of 23,147,854 regions. Regions were filtered for those located on autosomal chromosomes. MSIsensor *msi* was run on each BL and PD pair, ensuring that all regions had a minimum of 20X coverage and were located within SureSelect v5 target regions. All microsatellites that showed a significant difference in length distribution were manually reviewed to identify those that showed an increase in the PD sample. The ratio proportion of microsatellites with increased length variability divided by the total number of assessed microsatellites defines the MSI-score.

### *KRAS, NR AS* and *EGFR* mutation codon biases

Somatic mutation calls from The Cancer Genome Atlas (TCGA) were downloaded from the cBio web portal (Cerami et al., 2012; Gao et al., 2013) by selecting for ‘Colorectal Adenocarcinoma’ in the ‘PanCancer Atlas’. Mutation calls from studies (Bettegowda et al., 2014; Khan et al., 2018; Woolston et al., 2019) that reported the specific base change alterations in *KRAS, NRAS* and *EGFR* mutations in ctDNA were pooled to generate a comparative distribution from CRCs with acquired resistance to EGFR-AB. Only cases with KRAS/NRAS codon 12/13/61 mutations were included and these mutations were assessed. Mutation calls in *KRAS* were also identified from ctDNA in the CORRECT trial (Tabernero et al., 2015). Similarly, only KRAS codon 12/13/61 mutations were analyzed.

*EGFR* mutation calls in (Bettegowda et al., 2014; Woolston et al., 2019) were used to assess mutation codon biases in *EGFR* at acquired resistance.

### Kaplan-Meier Analysis

The *survfit* function in the Survival (v.2.44-1.1) R library was used to run the Kaplan-Meier analysis. Progression free survival (PFS) was measured from start of treatment to date of progression or death from any cause.

### Quantification And Statistical Analysis

All analyses were performed in R (v3.5.0) (Team, 2018). All p-values are two-sided and p<0.05 was considered significant. All t-tests were unpaired unless otherwise stated.

