## Supplementary material for "Mutational processes impact the evolution of anti-EGFR antibody resistance in colorectal cancer": Figure S1, Figure S2, Figure S3

#### Table of Contents

| | |
|---|---|
| FIGURE S1 | 2 |
| FIGURE S2 | 3 |
| FIGURE S3 | 4 |
| REFERENCES | 5 |

### Figure S1

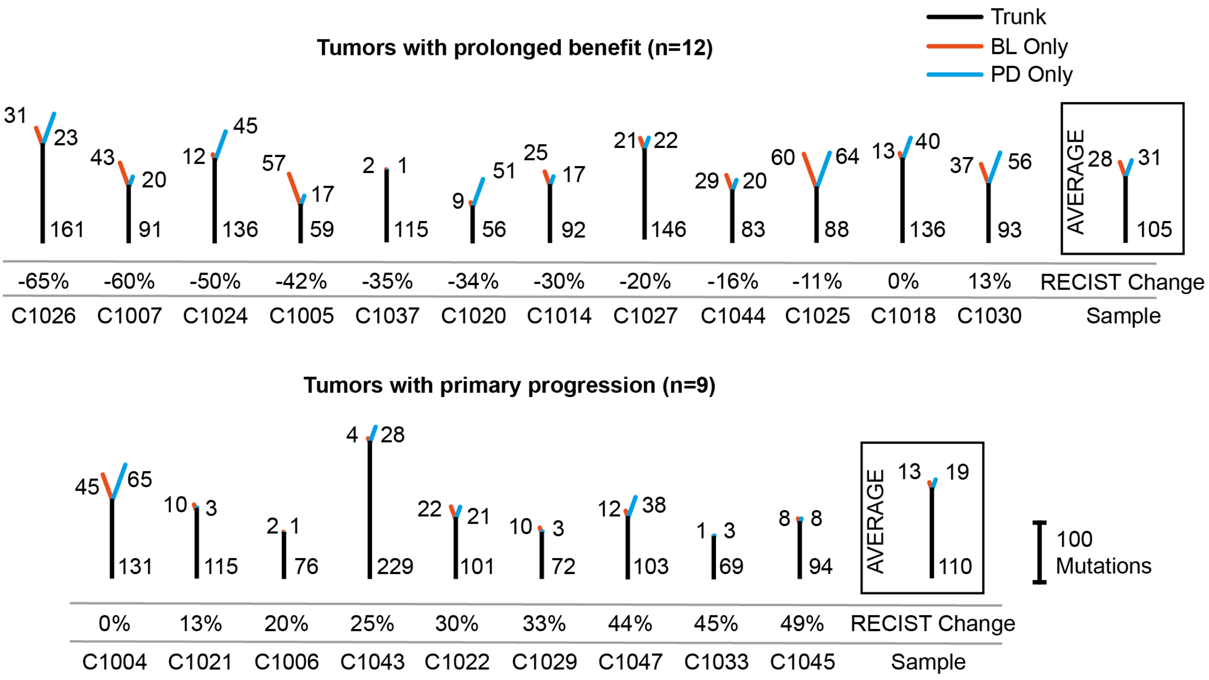


***Figure S1:*** *Clonal mutation trees for 21 tumors from the Prospect-C trial. Grouped into cases with prolonged benefit and primary progression. The numbers next to the trunk or the branches indicate clonal somatic mutations.*

**Figure S2**

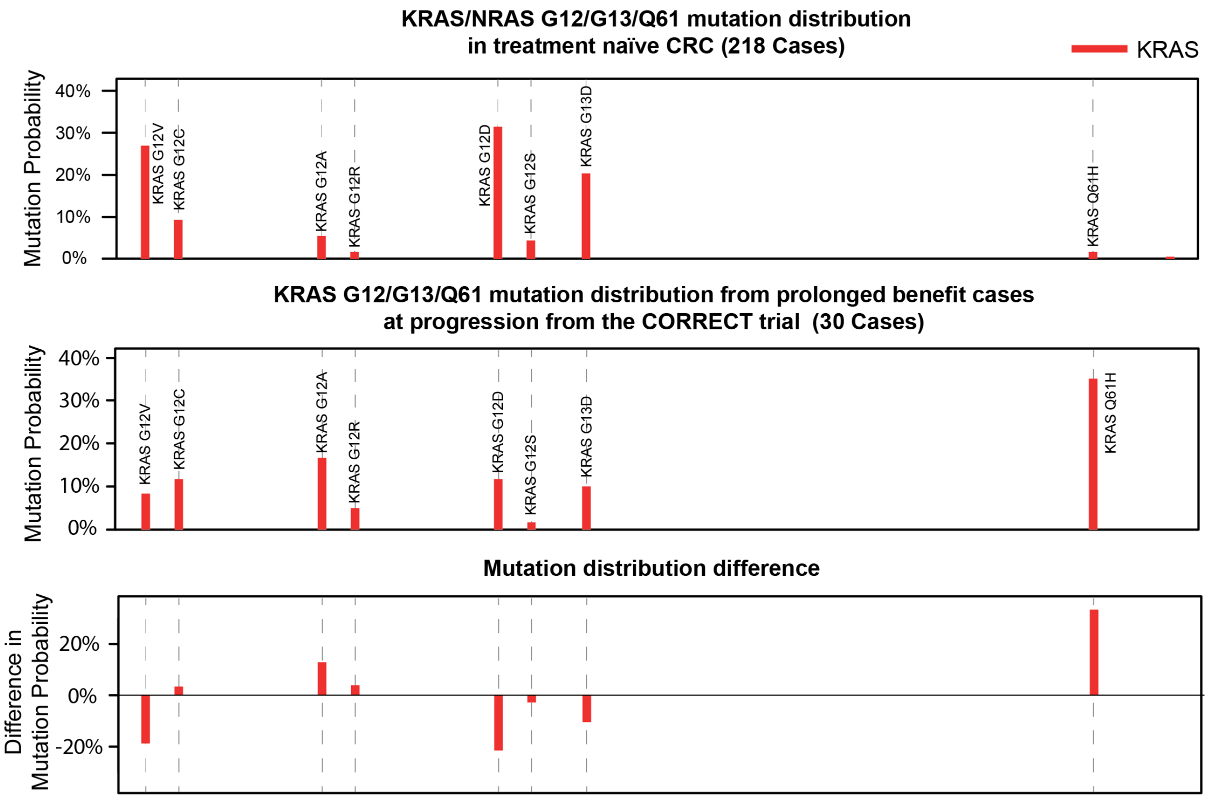


*Figure S2: Mutation frequency profiles of treatment naïve CRCs from the TCGA Pan-Cancer study versus the KRAS hotspot mutations identified in (Tabernero et al., 2015). The TCGA profile has been adjusted to only consider KRAS mutations that were assessed in the CORRECT trial.*

### Figure S3

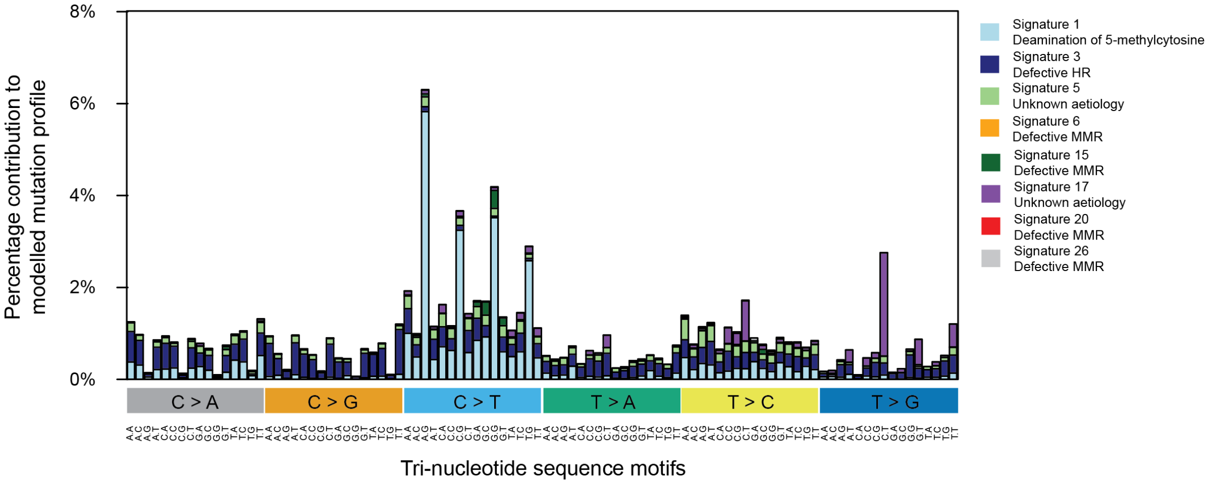


***Figure S3:*** *Modelled mutational profile based on the observed signature contributions at BL in tumors with prolonged benefit.*
